# Exome Sequencing Reveals a High Prevalence of *BRCA1* and *BRCA2* Founder Variants in a Diverse Population-Based Biobank

**DOI:** 10.1101/802348

**Authors:** Noura S. Abul-Husn, Emily R. Soper, Jacqueline A. Odgis, Sinead Cullina, Dean Bobo, Arden Moscati, Jessica E. Rodriguez, CBIPM Genomics Team, Regeneron Genetics Center, Ruth J.F. Loos, Judy H. Cho, Gillian M. Belbin, Sabrina A. Suckiel, Eimear E. Kenny

## Abstract

Pathogenic variants in *BRCA1* and *BRCA2* (*BRCA1/2*) lead to increased risk of breast, ovarian, and other cancers, but most variant positive individuals in the general population are unaware of their risk, and little is known about the prevalence of pathogenic *BRCA1/2* variants in non-European populations. We investigated *BRCA1/2* prevalence and impact using exome sequencing and electronic health record (EHR) data from 30,223 adult participants of the Bio*Me* Biobank in New York City. There were 218 (0.7%) individuals harboring expected pathogenic variants, resulting in an overall prevalence of 1 in 139. In sub-populations defined by genetic ancestry, the highest prevalence was in individuals of Ashkenazi Jewish (AJ; 1 in 49), Filipino and Southeast Asian (1 in 81), and Non-AJ European (1 in 103) descent. Among 218 variant positive individuals, 112 (51.4%) harbored known founder variants: 80 had AJ founder variants (*BRCA1* c.5266dupC and c.68_69delAG, and *BRCA2* c.5946delT), 7 had a Puerto Rican founder variant (*BRCA2* c.3922G>T), and 25 had one of 19 other founder variants. Non-European populations were more likely to harbor *BRCA1/2* variants that were not classified in ClinVar, or that had uncertain or conflicting evidence for pathogenicity. Within mixed ancestry populations, such as Hispanic/Latinos with genetic ancestry from Africa, Europe, and the Americas, there was a strong correlation between the proportion African genetic ancestry and the likelihood of harboring a *BRCA1/2* variant with uncertain or conflicting evidence for pathogenicity. Based on EHR and participant questionnaire data, ∼28% of variant positive individuals had a personal history, and ∼45% a personal or family history of *BRCA1/2*-associated cancers. Approximately 27% of variant positive individuals had evidence of prior clinical genetic testing for *BRCA1/2*. However, individuals with AJ founder variants were twice as likely to have had a clinical test (38%) than those with other pathogenic variants (19%). These findings deepen our knowledge about *BRCA1/2* variants and associated cancer risk in diverse populations, indicate a gap in knowledge about potential cancer-related variants in non-European populations, and suggest that genomic screening in diverse patient populations may be an effective tool to identify at-risk individuals.

## Background

The recognition of strong familial clustering of breast and ovarian cancer (1), followed by the discovery of the *BRCA1* and *BRCA2* (*BRCA1/2*) genes in 1994 (2) and 1995 (3), respectively, has led to the study and characterization of *BRCA1/2*-related hereditary breast and ovarian cancer syndrome (HBOC). Inherited pathogenic variants in either of these genes cause a significantly elevated risk for cancer of the female breast as well as high grade serous ovarian, tubal and peritoneal carcinoma. The risk for other cancers, including prostate, male breast, pancreas, melanoma and possibly others, is also increased (4). Pathogenic variants in these genes are highly penetrant and inherited in an autosomal dominant pattern.

The prevalence of pathogenic *BRCA1/2* variants has been previously estimated, with historical data suggesting a prevalence of approximately 1 in 400 individuals in the general population (5, 6). A higher prevalence has been observed in certain populations; for example, approximately 1 in 42 individuals of Ashkenazi Jewish (AJ) descent harbor one of three common founder variants (7, 8). Founder variants in other populations have also been described, including Icelandic, French Canadian, and Puerto Rican populations, and others (9). Recent unselected population-based genomic screening efforts have demonstrated a higher than expected prevalence of *BRCA1/2* pathogenic variants in predominantly European-ancestry individuals, approximately 1 in 190, with only half of these individuals meeting current guidelines for genetic testing (10-12) and only 18% having prior knowledge of their *BRCA1/2* status through clinical genetic testing (13).

Understanding of the prevalence and contribution to cancer risk of *BRCA1/2* variants in non-European populations has been limited by racial and ethnic disparities in genetic research (14). In addition to reduced uptake of genetic testing in diverse populations (15-18), there is a higher rate of detection of variants of uncertain significance in non-European populations (19-21). Here, we evaluated the range of *BRCA1/2* variants in a diverse patient population from the Bio*Me* Biobank in New York City, and explored clinical characteristics of individuals harboring expected pathogenic variants in *BRCA1/2*.

## Materials and Methods

### Setting and Study Population

The Bio*Me* Biobank is an electronic health record (EHR)-linked biobank of over 50,000 participants from the Mount Sinai Health System (MSHS) in New York, NY. Participant recruitment into Bio*Me* has been ongoing since 2007, and occurs predominantly through ambulatory care practices across the MSHS. The Bio*Me* participants in this analysis were recruited between 2007 and 2015, with approximately half coming from general medicine and primary care clinics and the rest from different specialty or multi-specialty sites at MSHS. Bio*Me* participants consent to provide DNA and plasma samples linked to their de-identified EHRs. Participants provide additional information on self-reported ancestry, personal and family medical history through questionnaires administered upon enrollment. This study was approved by the Icahn School of Medicine at Mount Sinai’s Institutional Review Board. The study population consisted of 30,223 consented Bio*Me* participants aged 18 years or older (upon enrollment), and with exome sequence data available through a collaboration with the Regeneron Genetics Center.

### Generation and QC of Genomic Data

Sample preparation and exome sequencing were performed at the Regeneron Genetics Center as previously described (22) yielding N=31,250 samples and n=8,761,478 sites. Genotype array data using the Illumina Global Screening Array was also generated for each individual (23). Post-hoc filtering of the sequence data included filtering of N=329 low-quality samples, including low coverage, contaminated and genotype-exome discordant samples; N=208 gender discordant and duplicate samples were also removed. This resulted in N=30,813 samples for downstream analysis, and N=30,223 samples from participants aged 18 years and older. Mean depth of coverage for remaining samples was 36.4x, and a minimum depth of 27.0x, and sequence coverage was sufficient to provide at least 20x haploid read depth at >85% of targeted bases in 96% of samples. Sites with missingness greater than 0.02 (n=267,955 sites) were removed, as were sites showing allele imbalance (n=320,877; allelic balance < 0.3 or > 0.8). Samples were stratified by self-reported ancestry, and sites with Hardy Weinberg equilibrium P<1×10^−6^ (n=12,762) were removed from analysis. Variants at multi-allelic sites in *<u>BRCA1</u>* and *BRCA2* (n=124) underwent the same quality control workflow as those from bi-allelic sites, with the exception that allelic balance was calculated only among heterozygous carriers of multi-allelic variants. Multi-allelic sites for which the mean allelic balance among heterozygous carriers was < 0.3 or > 0.8 were excluded from downstream analysis. This resulted in the exclusion of n=1 site, leaving a total of n=123 for further analysis. Manual inspection of pileups was performed for carriers (N=22) of the n=13 multi-allelic sites annotated as pathogenic in ClinVar. Of these, N=6 out of 7 carriers of the 13:32339421:C:CA variant were determined to be false positives and excluded from downstream analyses.

### Self-Reported and Genetic Ancestry

Self-reported ancestry categories were derived from a multiple-choice survey administered to participants upon enrollment into the Bio*Me* Biobank (23). Participants could select one or more of the following categories: African-American/African, American Indian/Native American, Caucasian/White, East/Southeast Asian, Hispanic/Latino, Jewish, Mediterranean, South Asian/Indian, or Other. Individuals who selected “Jewish”, “White/Caucasian”, or both were designated as “European American”. Individuals who selected “Mediterranean”, “Other”, or both were designated as “Other”. Individuals who selected multiple categories including “Hispanic/Latino” were designated as “Hispanic/Latino”. Individuals from the “Native American”, “Other”, or “Multiple Selected” categories were excluded from downstream analysis of prevalence in self-reported groups.

Genetic ancestry in the form of Identity-by-Descent community designation was performed on a subset of participants excluding second degree relatives and above, yielding 17 distinct communities representing patterns of cultural endogamy and recent diaspora to New York City. Eight of these communities with >400 unrelated participants were used for downstream analysis of prevalence. These communities included individuals with African-American and African ancestry (N=6,874), non-AJ European ancestry (N=5,474), AJ ancestry (N=3,887), Filipino and other Southeast Asian ancestry (N=556), as well as ancestry from Puerto Rico (PR; N=5,105), the Dominican Republic (DR; N=1,876), Ecuador (N=418), and other Central and South American communities (N=1,116). Full details of the global ancestry inference, genetic community detection, and genotype quality control are described in Belbin *et al*., 2019 (23). Finally, we determined the proportion African genetic ancestry in mixed ancestry Hispanic/Latino populations using the ADMIXTURE (24) software. We assumed five ancestral populations (k=5) with 5-fold cross validation across n=256,052 SNPs in N=27,984 unrelated participants that were also genotyped on the Global Screening Array (GSA), in addition to N=4,149 reference samples representing 5 continental regions (23). We estimated relatedness using the software KING (25), and for all prevalence estimates in self-reported and genetic ancestry groups, we excluded second degree relatives and above.

### BRCA1/2 Variant Annotation

Sequence variants were annotated with the Variant Effect Predictor (VEP; Genbank gene definitions; *BRCA1* NM_007294.3, *BRCA2* NM_000059.3). In order to reduce the set of false positive predicted loss-of-function (pLOF) calls, we also ran Loss-Of-Function Transcript Effect Estimator – LOFTEE, and defined the consensus calls from both methods as the set of pLOF variants for the study. Sequenced variants were cross-referenced with the ClinVar database (accessed July 2018) (26) and annotated according to their ClinVar assertions when available as pathogenic, likely pathogenic, uncertain significance, benign, likely benign, or with conflicting interpretations of pathogenicity. All variants with conflicting interpretations were manually reviewed in ClinVar (accessed November 2018) by a genetic counselor (J.A.O. or E.R.S.). In addition, we included the following categories of pLOF variants not classified in ClinVar: single nucleotide variants (SNVs) leading to a premature stop codon, loss of a start codon, or loss of a stop codon; SNVs or insertion/deletion sequence variants (indels) disrupting canonical splice acceptor or donor dinucleotides; open reading frame shifting indels leading to the formation of a premature stop codon. The union of ClinVar pathogenic/likely pathogenic and pLOF variants was termed “expected pathogenic”, and this set of variants was used to identify individuals in Bio*Me* for subsequent analyses of HBOC-related clinical characteristics.

### BRCA1/2 Founder Variants

All expected pathogenic variants detected in *BRCA1/2* were reviewed for evidence of a founder effect. This was carried out by manual review of each expected pathogenic variant by a genetic counselor (E.R.S.) in the Human Gene Mutation Database (27), ClinVar, and PubMed utilizing the currently designated HGVS nomenclature for each variant (28), as well as previous designations as noted in ClinVar. Variants were considered to be founder variants if they were described as such in the primary literature, based on confirmatory haplotype analysis or population frequency.

### Clinical Characteristics in Variant Positive Individuals

Individuals harboring expected pathogenic variants in *BRCA1/2* in Bio*Me*, termed “variant positive”, were evaluated for any evidence of personal or family histories of HBOC-related cancers, through extraction of International Classification of Diseases (ICD)-9 and ICD-10 codes from participant EHRs (**Supplementary Table S1**). These data were supplemented by participant questionnaire data for personal and family histories of HBOC-related cancers, which were available for 61 variant positive individuals. Medical record review of variant positive individuals was carried out independently by two individuals, including genetic counselors (J.A.O., E.R.S., or S.A.S.) and a clinical research coordinator (J.E.R.) to determine whether participants had evidence of previous clinical genetic testing for *BRCA1/2*. Data were summarized using medians and interquartile ranges (IQR) for continuous variables and frequencies and percentages for categorical variables. Pearson’s chi-squared test was used to test for statistical independence of different categorical outcomes measured in the study.

### HBOC-Related Cancers Case-Control and Phenome-Wide Association Studies

Cases were defined as participants having any of the ICD-9 or ICD-10 codes for personal history of HBOC-related cancers (**Supplementary Table S1**). Controls were defined as individuals without any of these ICD-9 or ICD-10 codes. We tested for association with variant positive compared to variant negative participants (defined as not having any variants that were pathogenic, uncertain/conflicting, or unclassified in ClinVar (novel)). Genotypes were coded using an additive model (0 for variant negative and 1 for variant positive). We repeated the analysis to compare participants with uncertain/conflicting variants with variant negative participants. We excluded individuals determined to be second degree relatives and above from the analysis. Odds ratios were estimated by logistic regression, and adjusted for age, sex and the first 5 principal components of ancestry.

We also performed a phenome-wide association study (PheWAS) of variant positive vs. variant negative participants using ICD-9 and ICD-10 based diagnosis codes that were collapsed to hierarchical clinical disease groups (termed phecodes) (29, 30). We performed logistic regression systematically using *BRCA1/2* expected pathogenic carrier status as the primary predictor variable and the presence of a given phecode as the outcome variable, excluding second degree relatives and above, and adjusting for age, sex and the first 5 principal components. To minimize spurious associations due to limited numbers of case observations, we restricted analyses to phecodes present at least 5 times among carriers, resulting in a total of p=260 tests. Statistical significance was determined using Bonferroni correction (Bonferroni adjusted significance threshold p<1.9×10^−4^). Logistic regression analyses were performed using PLINK (v1.90b3.35) software.

## Results

We evaluated *BRCA1/2* variants among 30,223 adult participants of the Bio*Me* Biobank with available exome sequence data and genotype array data. Participants were 59.3% female and had a median age of 59 years (**Table 1**). The majority of participants (74.3%) were of non-European descent, based on self-report. A total of 1601 variants were analyzed, including 1,478 (92.3%) occurring at bi-allelic sites and 123 (7.7%) at multi-allelic sites. The majority of variants were missense (63.5%), and 1,335 (83.4%) variants were available in ClinVar (**Supplementary Table S2**). The proportion of individuals harboring *BRCA1/2* variants that were not classified in ClinVar (novel) was lowest in individuals of self-reported European descent (0.8%), and highest in individuals of South Asian descent (2.3%; **Figure 1A**). The proportion of individuals harboring *BRCA1/2* variants of uncertain significance or with conflicting interpretations of pathogenicity (uncertain/conflicting) in ClinVar was lowest in individuals of self-reported European descent (4.1%) and highest in those of self-reported African/African-American descent (12.2%; **Figure 1B**). We saw a similar trend when investigating genetic ancestry within populations with recent mixed ancestry, for example Hispanic/Latino populations, who can trace their recent ancestry to Europe, Africa, and the Americas (**Supplementary Figure S1**). Although the mean uncertain/conflicting variant rate in all self-reported Hispanic/Latino participants was 8.5% (95% CI 7.9%-9.1%) (**Figure 1b**), this rate was almost two-fold higher in those with >60% African genetic ancestry (11.3% (95% CI 9.2%-13.9%) compared to those with <20% African genetic ancestry (6.9% (95% CI 6.1%-7.8%); chi-squared p<3.2×10^−4^; **Supplementary Figure S1**).

**Table 1.** Demographics of exome sequenced adult BioMe biobank participants, and of individuals harboring expected pathogenic variants in *BRCA1/2*.

|  | Sequenced BioMe participants (N = 30,223) | *BRCA1/2* variant negative (N = 20,760)* | *BRCA1/2* variant positive (N = 218) |
| --- | --- | --- | --- |
| Age, median (IQR) | 59 (45-70) | 59 (46-70) | 58 (43-70) |
| Female, N (%) | 17,914 (59.3) | 15,986 (59.1) | 137 (62.8) |
| Self-Reported Ancestry, N (%) |  |  |  |
| African/African-American | 6,878 (22.8) | 5,877 (28.3) | 33 (15.1) |
| East/Southeast Asian | 757 (2.5) | 659 (3.2) | 6 (2.8) |
| European | 7,772 (25.7) | 7,265 (35.0) | 121 (55.8) |
| Hispanic/Latino | 10,460 (34.6) | 9,360 (45.1) | 34 (15.6) |
| Native American | 52 (0.2) | 47 (0.2) | 0 (0) |
| South Asian | 605 (2.0) | 543 (2.6) | 0 (0) |
| Other | 2,343 (7.8) | 2,111 (10.2) | 13 (6.0) |
| Multiple selected | 1,125 (3.7) | 1,006 (4.9) | 10 (4.6) |
| Not Available | 231 (0.8) | 192 (0.9) | 1 (0.5) |
| *Variant negative participants are defined as not having any variants that were pathogenic, uncertain/conflicting, or unclassified in ClinVar |  |  |  |

**Figure 1.**
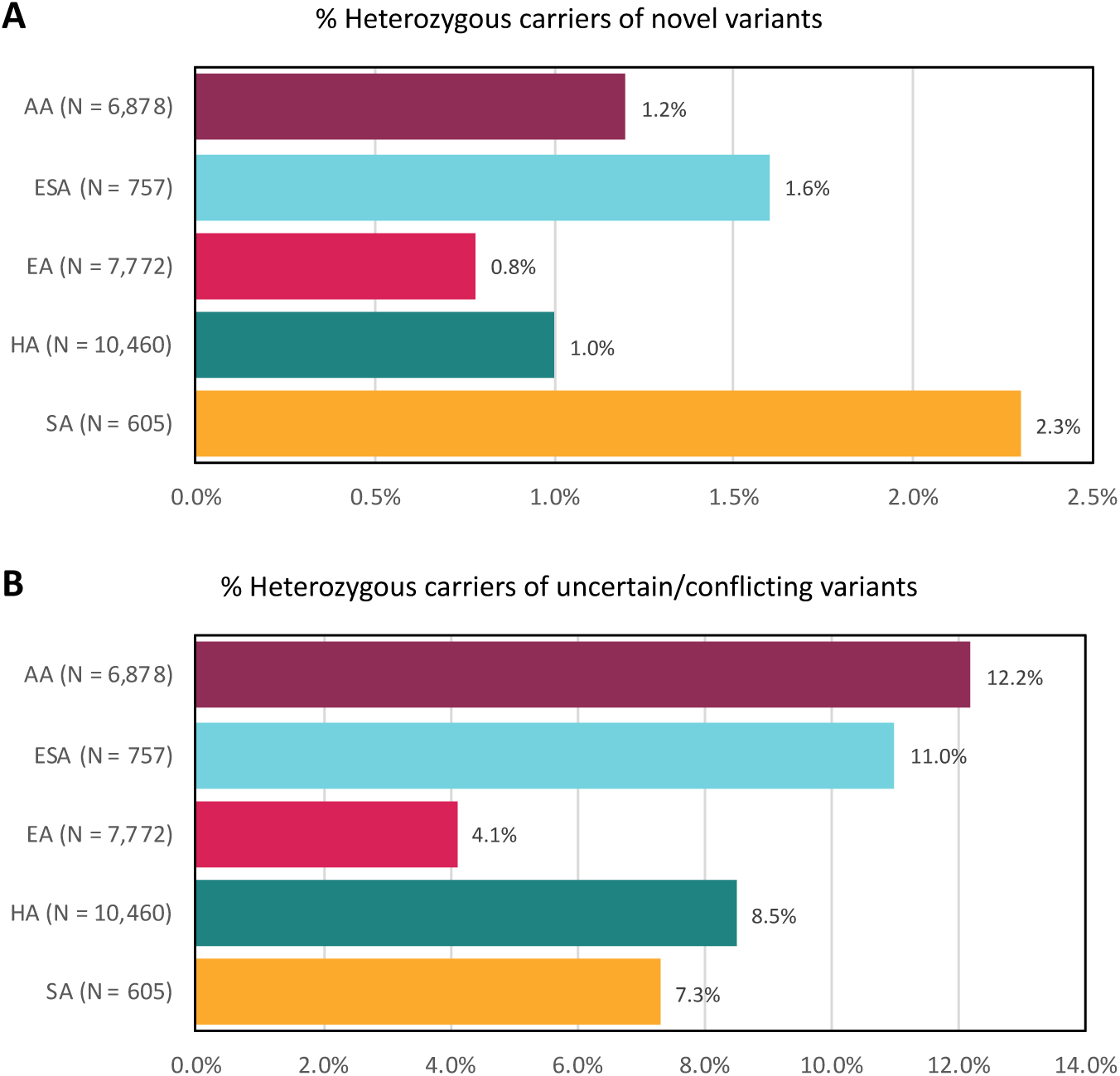
Among 1601 *BRCA1/2* variants identified in the Bio*Me* Biobank, there were 266 variants not classified in ClinVar (novel) and 635 variants of uncertain significance or with conflicting interpretations of pathogenicity in ClinVar (uncertain/conflicting). The proportion of individuals harboring novel (**Figure 1A**), or uncertain/conflicting (**Figure 1B**) variants varied across self-reported ancestry categories, and was lowest among individuals of European descent (0.8% and 4.1%, respectively). The proportion of individuals harboring novel variants was highest in individuals of South Asian descent (2.3%), and the proportion harboring uncertain/conflicting variants was highest in individuals of African/African-American descent (12.2%). AA, African/African-American descent; ESA, East/Southeast Asian descent; EA, European descent; HA, Hispanic/Latino descent; SA, South Asian descent.

Exome sequence data of the *BRCA1/2* genes was then used to identify expected pathogenic variants. There were 102 variants with a pathogenic or likely pathogenic assertion in ClinVar, all of which had a 2- or 3-star review status (**Supplementary Table S3**). There were 10 additional pLOF variants (frameshift or stop gained) that were not classified in ClinVar, including 2 in *BRCA1* and 8 in *BRCA2*. The 10 pLOF variants were each observed as singletons in Bio*Me*, and only one of them (*BRCA2* c.1039C>T) was found in the gnomAD database (31) with an allele frequency of 0.000004, suggesting that these are rare in the general population. The union of 102 ClinVar pathogenic and 10 additional rare pLOF variants was the set of expected pathogenic *BRCA1/2* variants (N = 112) used to define variant positive individuals in Bio*Me*.

Overall, 218 (0.7%) individuals in Bio*Me* harbored expected pathogenic variants in *BRCA1/2*: 86 (39.4%) of these individuals had an expected pathogenic variant in *BRCA1*, 131 (60.1%) had a variant in *BRCA2*, and 1 (0.5%) individual had a variant in both *BRCA1* (c.68_69delAG) and *BRCA2* (c.5946delT). Variant positive individuals were 62.8% female and had a median age of 58 years (**Table 1**). The prevalence of Bio*Me* participants harboring expected pathogenic variants in *BRCA1/2* was 1:139 (**Table 2**). In a subset of individuals excluding second degree relatives and above (N=27,816), overall prevalence was unchanged at 1:134. In the unrelated subset, prevalence was highest in individuals of self-reported European descent (1:66) and lowest in those of Hispanic/Latino descent (1:283). We previously used genotype array data to identify fine-scale population groups in Bio*Me* using genetic ancestry (23), revealing eight communities with greater than 400 individuals represented (**Table 2**). Across these, prevalence was highest in individuals with AJ ancestry (1:49), among whom the majority (72 out of 80 individuals, or 90.0%) harbored one of the three AJ founder variants (c.5266dupC and c.68_69delAG in *BRCA1*, and c.5946delT in *BRCA2*), and 8 individuals (10.0%) harbored a different variant in *BRCA1/2* (**Supplementary Table S3**). Prevalence was lower in non-AJ Europeans (1:103), and lowest in those with ancestry from PR (1:341) and DR (1:469; **Table 2**).

**Table 2.** Prevalence of expected pathogenic *BRCA1/2* variants in the Bio*Me* Biobank. We assessed the prevalence of *BRCA1/2* variants in all sequenced participants, in an unrelated subset of participants, across self-reported ancestry groups, and across genetic ancestry groups for which there were greater than 400 individuals.

| Population Characteristics | N | *BRCA1/2* Variant Positive, N (%) | Estimated Prevalence |
| --- | --- | --- | --- |
| All sequenced participants | 30,223 | 218 (0.7) | 1:139 |
| **Unrelated subset** - including only one individual in every first- and second-degree relationship | 27,816 | 208 (0.7) | 1:134 |
| **Self-reported ancestry (unrelated subset)** |  |  |  |
| African/African-American | 6,236 | 31 (0.5) | 1:201 |
| East/Southeast Asian | 739 | 6 (0.8) | 1:123 |
| European | 7,600 | 116 (1.5) | 1:66 |
| Hispanic/Latino | 9,050 | 32 (0.4) | 1:283 |
| Native American | 47 | 0 (0) | - |
| South Asian | 585 | 0 (0) | - |
| Other | 2,271 | 13 (0.6) | - |
| Multiple selected | 1,078 | 9 (0.7) | - |
| Not Available | 211 | 1 (0.5) | - |
| **Genetic ancestry (unrelated subset)** |  |  |  |
| African-American and African | 6,874 | 31 (0.5) | 1:222 |
| Ashkenazi Jewish | 3,889 | 80 (2.1) | 1:49 |
| Non-Ashkenazi Jewish European | 5,474 | 53 (1.0) | 1:103 |
| Filipino and Southeast Asian | 566 | 7 (1.2) | 1:81 |
| Dominican | 1,876 | 4 (0.2) | 1:469 |
| Ecuadorian | 418 | 2 (0.5) | 1:209 |
| Puerto Rican | 5,105 | 15 (0.3) | 1:365 |
| Other Central and South American | 1,116 | 8 (0.7) | 1:140 |

We identified 23 unique founder variants that have previously been reported in multiple founder populations, including 13 variants in *BRCA1* and 10 in *BRCA2* (**Table 3**). A total of 112 of 218 variant positive individuals (51.4%) were identified as harboring at least one founder variant (61 individuals with a variant in *BRCA1*, 50 with *BRCA2*, and 1 with both *BRCA1* and *BRCA2*). The majority of identified founder variants were accounted for by the three AJ founder variants, with 80 individuals in Bio*Me* harboring at least one of these variants, 72 of whom had AJ genetic ancestry. There were 32 participants harboring non-AJ founder variants in *BRCA1/2*, the most common being *BRCA2* c.3922G>T, a well-documented founder variant in PR (32). Among 15 *BRCA1/2* variant positive individuals with genetic ancestry from PR, 7 (46.7%) harbored the *BRCA2* c.3922G>T variant, and 3 others (20.0%) harbored Chilean or Spanish founder variants (**Table 3**).

**Table 3.**
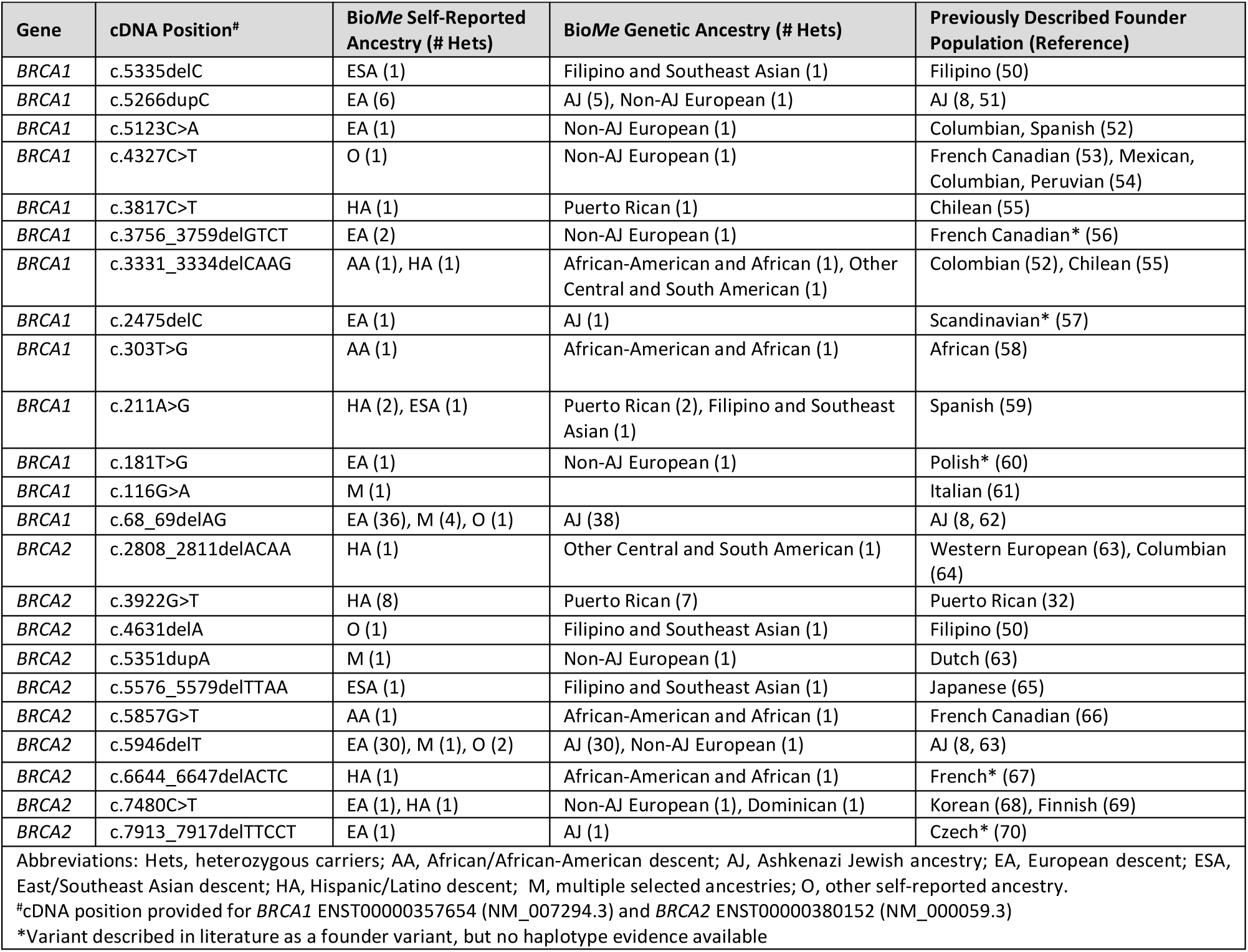
Founder variants identified among 112 *BRCA1/2* expected pathogenic variants in the Bio*Me* Biobank.

We evaluated the clinical characteristics of *BRCA1/2* variant positive individuals using EHR-extracted diagnosis codes (**Supplementary Table S1**), as well as additional personal and family medical history questionnaire data available for 61 of these individuals. Overall, 61 of 218 (28.0%) *BRCA1/2* variant positive individuals had a documented personal history and 98 (45.0%) had either a personal or family history of HBOC-related cancer (breast, ovarian, pancreatic, prostate, or melanoma; **Table 4**). Variant positive females were 2.8 times more likely than males to have a personal or family history of HBOC-related cancers (chi-squared p<1.0×10^−6^). Among variant positive females (N = 137), 53 (38.7%) had HBOC-related cancers, including 50 (36.5%) with breast or ovarian cancer. Among the three females with cancer other than breast or ovarian, two had pancreatic cancer and one had melanoma. There were 3 (2.2%) variant positive females who had more than one cancer, all of whom had both breast and ovarian cancer: one with *BRCA1* c.68_69delAG, and two with *BRCA2* c.5946delT. Among variant positive males (N = 81), 2 (2.5%) had breast cancer (*BRCA1* c.5266dupC and *BRCA2* c.4471_4474delCTGA), and 6 (7.4%) had prostate cancer (two men with *BRCA1* c.5266dupC, and one man each with *BRCA1* c.68_69delAG, *BRCA2* c.2808_2811delACAA, *BRCA2* c.5946delT, and *BRCA2* c.4716_4717delinsAAAGACC). One of these men (1.2%) had more than one cancer (breast and pancreatic) and harbored *BRCA2* c.4471_4474delCTGA.

**Table 4.**
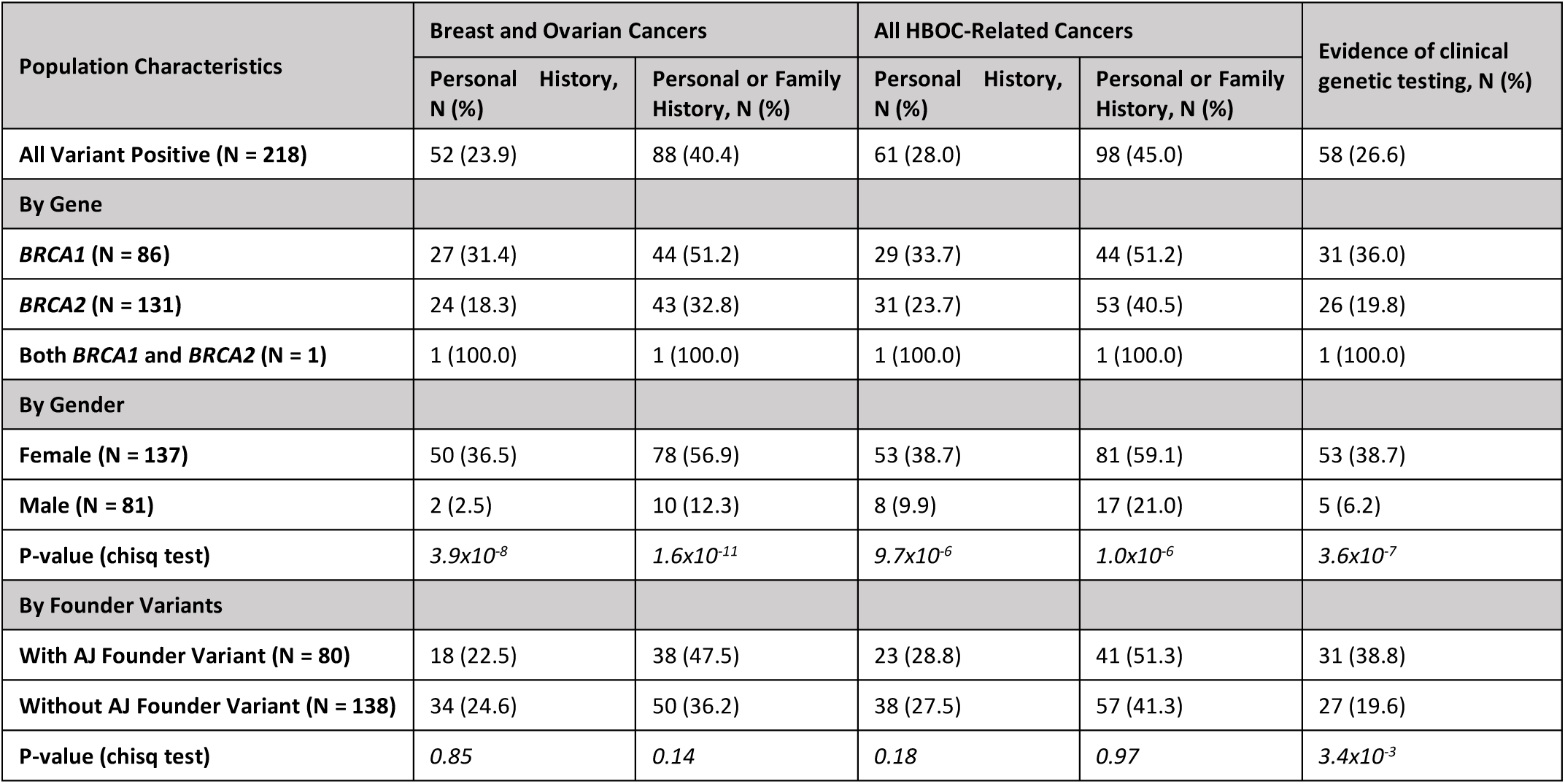
Clinical characteristics of *BRCA1/2* variant positive individuals. Evidence of HBOC-related cancers (breast, ovarian, prostate, pancreatic, and melanoma) and of clinical genetic testing among 218 Bio*Me* biobank participants harboring expected pathogenic *BRCA1/2* variants.

We assessed the number of variant positive individuals with prior knowledge of their *BRCA1/2* variant status. Review of medical records revealed that 58 (26.6%) had EHR evidence of clinical genetic testing for *BRCA1/2* (**Table 4**). Among 98 variant positive individuals with a personal or family history of HBOC-related cancer, 51 (52.0%) had evidence of clinical genetic testing. Only 5 of 81 (6.2%) males had evidence of clinical genetic testing, compared to 53 of 137 (38.7%) females (chi-squared p< 3.6×10^−7^). Although personal rates of cancer were similar among individuals with AJ founder variants and those with other variants (28.8% vs. 27.5%, chi-squared p=0.18), knowledge of *BRCA1/2* variant status varied: 31 of 80 (38.8%) individuals with AJ founder variants had documented evidence of clinical genetic testing, compared to only 27 of 138 (19.6%) individuals harboring other *BRCA1/2* variants (chi-squared p<3.4×10^−3^).

We tested for association with HBOC-related cancers in variant positive (N=208) compared to variant negative (not harboring any ClinVar pathogenic, uncertain/conflicting, or novel variants; N=24,927) participants in the unrelated subset. Variant positive individuals had increased odds of HBOC-related cancers (odds ratio (OR) 5.6; 95% confidence interval (CI) 4.0 to 8.0; p=6.7×10^−23^). In contrast, participants harboring uncertain/conflicting variants (N=2,395) did not have increased odds of HBOC-related cancers (OR 1.2; 95% CI 1.0 to 1.4; p=0.1). To more comprehensively evaluate the clinical consequences of expected pathogenic variants in *BRCA1/2*, we performed a PheWAS of variant positive vs. variant negative participants. Using a Bonferroni significance threshold of p=1.9×10^−4^ for associations with 260 clinical diagnoses, we identified significant associations with “Malignant neoplasm of female breast” (OR 8.1; 95% CI 5.4 to 12.2; p=2.2×10^−23^) and “Other specified disorders of breast” (OR 6.9; 95% CI 2.9 to 16.2; p=9.0x^10-6^; **Supplementary Figure S2**). There were no associations with other types of cancer or non-cancer phenotypes, including known HBOC-related cancers, suggesting we may have been underpowered to observe other relevant associations.

## Discussion

In this study, we demonstrate the ability of large-scale, population-based genomic sequencing to identify and characterize consequential variants in *BRCA1/2* in a large, ethnically diverse health system. We found an overall prevalence of 1 in 139 individuals with expected pathogenic variants in *BRCA1/2*, observed differing frequencies of such variants among a broad range of represented ancestries, and discovered that the majority of individuals harboring these variants were unaware of their genomic risk status.

The overall prevalence of expected pathogenic *BRCA1/2* variants in our population was higher than previous estimates (5, 6, 13), and may be partly explained by the large number of founder variants detected. The highest prevalence was 1 in 49 (2.1%) in individuals with AJ genetic ancestry, which is similar to the previously established prevalence of 1 in 42 (2.3%) in this population (7, 8). The high proportion of AJ individuals in our cohort (14.0%) contributed to the high overall prevalence observed. Multiple other founder variants were also detected in different populations in our study, including the c.3922G>T (p.Glu1308Ter) variant in *BRCA2* that we found in almost half of the variant positive individuals with ancestry from PR, consistent with previous findings (32). We report, for the first time, prevalence estimates in a number of diverse populations, including African American and Hispanic/Latino populations for which these estimates did not previously exist.

Our findings also revealed that non-European populations, and particularly those most genetically divergent from European populations, are more likely to harbor *BRCA1/2* variants that are not classified in public databases, or that have uncertain or conflicting evidence for pathogenicity. This was also evident in mixed ancestry populations such as Hispanic/Latino populations, in whom the proportion of variants with uncertain/conflicting interpretations correlated with the percent African genetic ancestry. While *BRCA1/2* variant positive individuals had significantly increased risk of HBOC-related cancers, those with uncertain/conflicting variants did not, suggesting that many of these variants are likely to be benign or of low penetrance. These data add to a growing body of literature (19-21) underscoring the pressing need to further characterize genomic variation across diverse populations.

As with previous studies, there was a higher rate of relevant cancers in *BRCA1* variant positive individuals than in *BRCA2*, and in women than in men (13, 33, 34). Over one-third of the variant positive females in our study had a documented current or prior diagnosis of a HBOC-related cancer. Genomic screening in individuals with cancer still provides an opportunity for early detection or prophylaxis, as evidenced by the finding of a second primary cancer in four participants. Genomic screening in apparently healthy men may represent an opportunity for intervention through increased prostate surveillance, given the recently recognized contribution of germline *BRCA1*/2 variants to metastatic prostate cancer burden (35).

Knowledge of *BRCA1/2* status as documented in participant EHRs was only 27% overall, and even lower (20%) in individuals with non-AJ founder variants, confirming prior reports of clinical under-ascertainment (13). Of note, 10% of the variant positive AJ individuals harbored non-founder variants, consistent with previous findings (36), and highlighting the need for comprehensive testing of *BRCA1/2* genes rather than targeted screening for specific founder variants in this population. The observed difference in clinical testing among individuals with or without AJ founder variants, despite similar rates of cancer, indicates that there may be additional barriers to genetic testing in populations that are not considered higher risk on the basis of ancestry. Obstacles in non-AJ populations could include lack of patient awareness about *BRCA1/2*, lower suspicion for HBOC by healthcare providers, or reduced access and/or uptake of genetic testing in certain populations within the context of broader healthcare disparities. Such barriers have been described in African American and Hispanic/Latino populations, the two largest non-European populations in Bio*Me*, suggesting that interventions to improve awareness, risk-perception, and patient-provider communication are needed to reduce disparities in *BRCA1/2* testing in diverse populations (37).

Current evidence- and expert opinion-driven guidelines (10, 11, 38) as well as statistical models (39-42) to identify potential candidates for *BRCA1/2* testing are mainly based on the number of individuals with relevant cancers in a kindred, age(s) of diagnosis, and ancestry. Testing criteria have widened over time with the recognition that they do not sufficiently identify all individuals harboring a *BRCA1/2* pathogenic variant. Nevertheless, our findings suggest that current clinical practices still miss a significant opportunity for reducing morbidity and mortality through identification of high-risk variant positive individuals. While we were unable to evaluate whether variant positive individuals would meet current testing criteria, we did observe that almost half of those with a relevant personal or family history of cancer had no evidence of clinical *BRCA1/2* testing. The potential for improved health outcomes from genomic screening through ascertainment of patients and identification of at-risk relatives through cascade testing (43, 44) supports the Centers for Disease Control and Prevention’s designation of HBOC as a tier 1 genomic condition for which positive public health impact exists (https://www.cdc.gov/genomics/implementation/toolkit/tier1.htm).

There are limitations to our study. The study population consisted of individuals recruited from clinical care sites, which does not necessarily reflect the general population of New York City. However, these findings do provide insight into diverse patient populations that were ascertained in a relatively unselected, population-based manner, and that have not been previously represented in similar research efforts. The observed prevalence of *BRCA1/2* expected pathogenic variants may represent an underestimate, as certain variants would not be detected via this approach, including large copy number variants, which make up approximately 10% of all *BRCA1/2* pathogenic variants (45-48). Additionally, some percentage of variants of uncertain significance may in fact be pathogenic, and likely will be classified as such in the future. We were also constrained by the use of EHR-extracted clinical information, which may not reflect complete medical and family history (49), and may downwardly bias the true penetrance of HBOC in our cohort.

## Conclusions

Genomic screening for pathogenic *BRCA1/2* variants in apparently healthy individuals has the potential to lead to earlier diagnosis of cancer via increased surveillance, as well as cancer risk reduction via prophylactic medical interventions. In this study, we provide evidence for a higher overall prevalence of *BRCA1/2* expected pathogenic variants in the Bio*Me* Biobank than historically appreciated, in line with recent findings from another unselected clinical care cohort (13). We show that this approach can effectively identify at-risk individuals across ethnically diverse and underserved populations such as those present in Bio*Me*. These findings are in part due to the cross-sectional representation of founder variants from multiple different populations, which accounted for over half of individuals harboring pathogenic variants in this study. We demonstrate that genomic screening for *BRCA1/2* in diverse patient populations may be an effective tool to identify otherwise unrecognized HBOC-associated variants, in order to prevent or diagnose disease. However, further work is needed to accurately classify pathogenic variants in non-European populations, in order to most effectively use this strategy to improve health outcomes in diverse settings.

## Declarations

### Ethics Approval and Consent to Participate

This study was approved by the Icahn School of Medicine at Mount Sinai’s Institutional Review Board. The study population consisted of 30,223 consented participants aged 18 years or older from Mount Sinai’s Bio*Me* Biobank.

### Availability of Data and Materials

Expected pathogenic variants in *BRCA1*/*2* reported in this paper are tabulated in Supplementary Table S3. The exome sequencing datasets generated and/or analyzed during the current study are not publicly available, but summary statistics are available from the corresponding author on reasonable request.

### Authors’ Contributions

G.M.B, S.C., A.M., and D.B. performed sequence data QC, annotation, and analysis. G.M.B. and S.C. performed genetic ancestry analysis. N.S.A-H., J.A.O., E.R.S., and S.A.S. analyzed and interpreted *BRCA1/2* sequence data. N.S.A-H., E.R.S., and J.A.O. analyzed and interpreted EHR data. E.R.S., S.A.S., J.A.O. and J.E.R. reviewed medical records for evidence of clinical genetic testing. N.S.A-H., E.R.S., J.A.O., G.M.B, and E.E.K. contributed to writing the manuscript. N.S.A-H. and E.E.K. designed the study and supervised all aspects of the analysis and manuscript preparation. All authors read and approved the final manuscript.

## Acknowledgements

We would like to thank participants of the Bio*Me* Biobank for their permission to use their health and genomic information.

