## Supplementary Information for "Exome Sequencing Reveals a High Prevalence of *BRCA1* and *BRCA2* Founder Variants in a Diverse Population-Based Biobank"

All authors/contributors are listed in alphabetical order.

**CBIPM Leadership Team**
Noura Abul-Husn, Erwin Bottinger, Judy Cho, Ron Do, Steve Ellis, Omri Gottesman, Yuval Itan, Eimear Kenny, Ruth Loos, Amanda Merkelson, Girish Nadkarni, Aniwaa Owusu-Obeng.

Contribution: All authors contributed to securing funding, study design and oversight.

**Sequencing and Lab Operations**
Bernadette Liggayu, Amanda Merkelson, Janice Morinigo, Patrick Shanley, Quingbin Song

Contribution: All authors are responsible for DNA extraction, sample handling and tracking, and the library information management system.

**Clinical Informatics**
Noura Abul-Husn, Lili Chan, Steve Ellis, Omri Gottesman, Arden Moscati, Girish Nadkarni, Rajiv Nadukuru, Aniwaa Owusu-Obeng, Teilman Van Vlick

Contribution: All authors are responsible for analysis needed to produce electronic health record extracted data.

**Genome Informatics**
Gillian Belbin, Dean Bobo, Kumardeep Chaudhary, Sinead Cullina, Ron Do, Aine Duffy, Amanda Dobbyn, Yuval Itan, Eimear Kenny, Margret Linan, Carla Marquez-luna, Arden Moscati, Ha My, Micheal Preuss, Cigdem Sevim, Stephane Wenric, Ryan Walker, Zhe Wang, Yiming Wu.

Contribution: All authors are responsible for analysis needed to produce exome and genotype data.

**Regeneron Genetics Center Banner Author List and Contribution Statements**

All authors/contributors are listed in alphabetical order.

**RGC Management and Leadership Team**

Goncalo Abecasis, Aris Baras, Michael Cantor, Giovanni Coppola, Aris Economides, John D. Overton, Jeffrey G. Reid, Alan Shuldiner.

Contribution: All authors contributed to securing funding, study design and oversight. All authors reviewed the final version of the manuscript.

**Sequencing and Lab Operations**

Christina Beechert, Caitlin Forsythe, Erin D. Fuller, Zhenhua Gu, Michael Lattari, Alexander Lopez, John D. Overton, Thomas D. Schleicher, Maria Sotiropoulos Padilla, Karina Toledo, Louis Widom, Sarah E. Wolf, Manasi Pradhan, Kia Manoochehri, Ricardo H. Ulloa.

Contribution: C.B., C.F., K.T., A.L., and J.D.O. performed and are responsible for sample genotyping. C.B, C.F., E.D.F., M.L., M.S.P., K.T., L.W., S.E.W., A.L., and J.D.O. performed and are responsible for exome sequencing. T.D.S., Z.G., A.L., and J.D.O. conceived and are responsible for laboratory automation. M.P., K.M., R.U., and J.D.O are responsible for sample tracking and the library information management system.

**Genome Informatics**

Xiaodong Bai, Suganthi Balasubramanian, Leland Barnard, Andrew Blumenfeld, Yating Chai, Gisu Eom, Lukas Habegger, Young Hahn, Alicia Hawes, Shareef Khalid, Jeffrey G. Reid, Evan K. Maxwell, John Penn, Jeffrey C. Staples, Ashish Yadav.

Contribution: X.B., A.H., Y.C., J.P., and J.G.R. performed and are responsible for analysis needed to produce exome and genotype data. G.E., Y.H., and J.G.R. provided compute infrastructure development and operational support. S.K., S.B., and J.G.R. provide variant and gene annotations and their functional interpretation of variants. E.M., L.B., J.S., A.B., A.Y., L.H., J.G.R. conceived and are responsible for creating, developing, and deploying analysis platforms and computational methods for analyzing genomic data.

**Planning, Strategy, and Operations**

Paloma M. Guzzardo, Marcus B. Jones, Lyndon J. Mitnau.

Contribution: All authors contributed to the management and coordination of all research activities, planning and execution. All authors contributed to the review process for the final version of the manuscript.

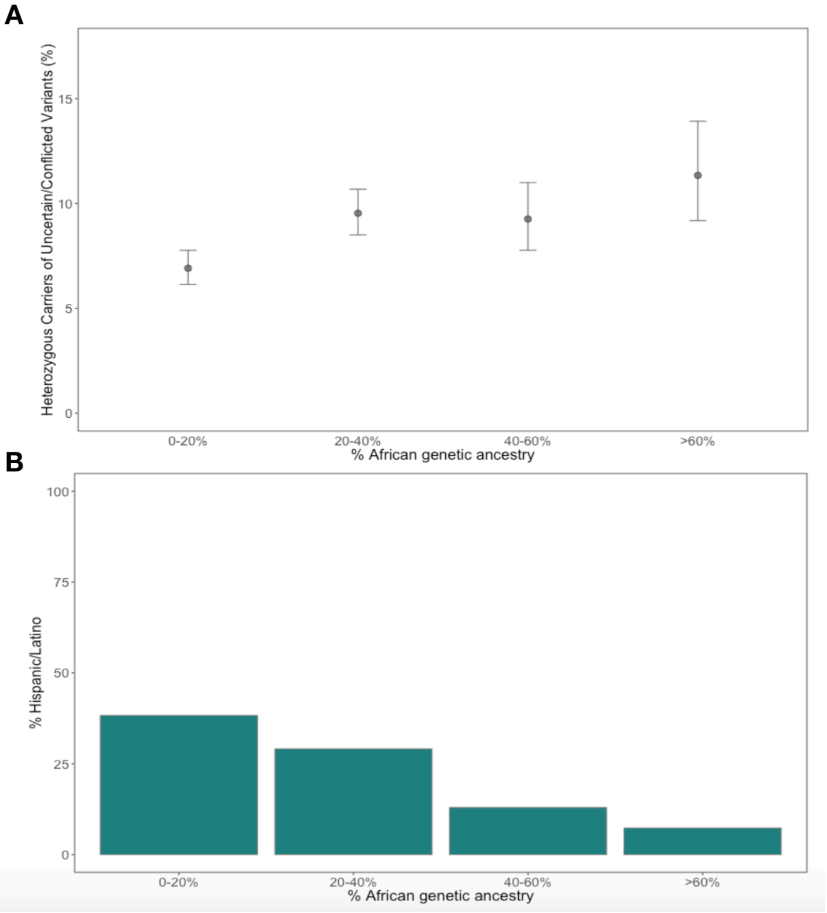


**Supplementary Figure S1.** Among 394 *BRCA1/2* variants identified in unrelated, self-reported Hispanic/Latinos (N=10,460) in the Bio*Me* Biobank, there were 125 variants of uncertain significance or with conflicting interpretations of pathogenicity in ClinVar (uncertain/conflicting). We examined the proportion of uncertain/conflicting variants in four bins of African genetic ancestry (0%-20%, 21%-40%, 41%-60%, 61%-100%) and found a strong correlation (Pearson correlation coefficient 0.96) between higher proportion African genetic ancestry and higher rates of uncertain/conflicting variants (**1A**). Hispanic/Latino participants in the highest African genetic ancestry bin had an almost two-fold higher rate of uncertain/conflicting variants (11.3%, 95% CI 9.2%-13.9%) compared to those in the lowest bin (6.9%, 95% CI 6.1%-7.7%); chi-squared test $p<3.2x10^{-4}$**. 1B** shows the breakdown of self-reported Hispanic/Latinos by African genetic ancestry bins; 0-20% (N=4,007); 21%-40% (N=3,044); 41%-60% (N=1,357); 61%-100% (N=766).

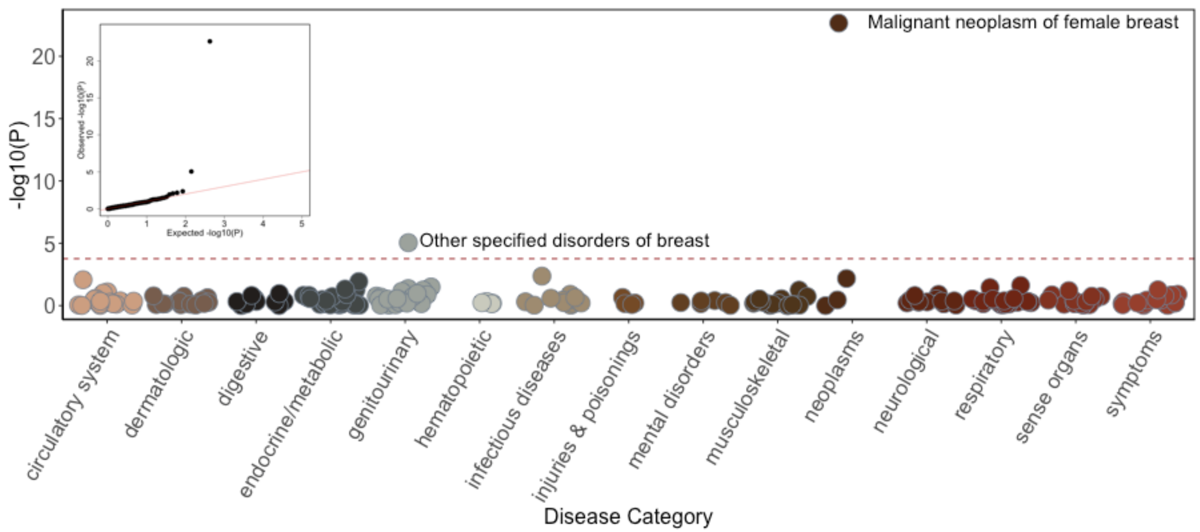


**Supplementary Figure S2.** Phenome-wide association study of variant positive vs. variant negative participants using EHR-derived clinical diagnoses. We restricted analyses to diagnoses present at least 5 times among variant positive participants, resulting in a total of 260 tests. Logistic regression was performed adjusting for age, sex, and the first 5 principal components. Using a Bonferroni adjusted significance threshold of $p<1.9x10^{-4}$, we identified two significant associations of variant positive participants with "Malignant neoplasm of female breast" (OR 8.1; 95% CI 5.4 to 12.2; $p=2.2x10^{-23}$) and "Other specified disorders of breast" (OR 6.9; 95% CI 2.9 to 16.2; $p=9.0x^{10-6}$).

**Supplementary Table S1.** International Classification of Diseases (ICD)-9 and -10 codes used to characterize *BRCA1/2* variant positive individuals.

| Personal and Family Medical History | | ICD-9 Codes | ICD-10 Codes | Definition |
|---|---|---|---|---|
| **Personal History** | Breast cancer | 174.*, 175.*, 233.0, V10.3 | C50.*, D05.*, Z85.3, Z86.000 | Current or past diagnosis of invasive and/or in situ carcinoma of the breast (male or female) |
| | Ovarian cancer | 183.*, V10.43 | C56.*, C57.0*, Z85.43 | Current or past diagnosis of carcinoma of the ovary |
| | Prostate cancer | 185.*, 233.4, V10.46 | C61, Z85.46, D07.5 | Current or past diagnosis of carcinoma of the prostate |
| | Pancreatic cancer | 157.* | C25.*, Z85.07 | Current or past diagnosis of carcinoma of the pancreas |
| | Melanoma | 172.*, V10.82 | C43*, Z85.820 | Current or past diagnosis of cutaneous melanoma |
| | Prior Genetic Testing | V84.01, V84.02, V84.03 | Z15.01, Z15.02, Z15.03 | Genetic susceptibility to breast, ovary or prostate |
| **Family History** | Breast cancer | V16.3 | Z80.3 | Family history of breast cancer |
| | Ovarian cancer | V16.41 | Z80.41 | Family history of ovarian cancer |
| | Prostate cancer | V16.42 | Z80.42 | Family history of prostate cancer |
| | Gastrointestinal tract cancer | V16.0 | Z80.0 | Family history of cancer of the gastrointestinal tract (including pancreas) |
| | Prior Genetic Testing | V18.9 | Z84.81 | Family history of genetic disease |

**Supplementary Table S2.** Distribution of 1601 *BRCA1/2* variants obtained from exome sequence data available from 30,223 adult Bio*Me* Biobank participants, according to ClinVar assertion and variant type.

| **ClinVar Assertion** | ***BRCA1*, N (%)** | ***BRCA2*, N (%)** | ***BRCA1/2*, N (%)** |
|---|---|---|---|
| Pathogenic/Likely Pathogenic | 31 (5.6) | 71 (6.8) | 102 (6.4) |
| Uncertain Significance | 174 (31.4) | 356 (34.0) | 530 (33.1) |
| Conflicting Interpretations | 27 (4.9) | 78 (7.5) | 105 (6.6) |
| Benign/Likely Benign | 238 (42.9) | 360 (34.4) | 598 (37.4) |
| NA | 85 (15.3) | 181 (17.3) | 266 (16.6)* |
| **Variant Type#** | | | |
| Frameshift | 12 (2.2) | 44 (4.2) | 56 (3.5) |
| Missense | 361 (65.0) | 655 (62.6) | 1016 (63.5) |
| Stop gained | 12 (2.2) | 19 (1.8) | 31 (1.9) |
| Synonymous | 150 (27.0) | 221 (21.1) | 371 (23.2) |
| NA | 20 (3.6) | 107 (10.2) | 127 (7.9) |
| **Total** | **555** | **1046** | **1601** |

*10 of the 266 variants with no ClinVar assertions were predicted loss-of-function variants (frameshift or stop gained) and were included in the set of expected pathogenic variants used to define variant positive individuals.

#Consequence based on Variant Effect Predictor annotation

**Supplementary Table S3.** BRCA1/2 expected pathogenic variants identified in 30,223 exome sequenced adults from the Bio*Me* Biobank.

| CHR:POS:REF:ALT | Gene | Function | rsID | cDNA Position | Protein Position | ClinVar Significance (Review Status) | # Hets | Self-Reported Ancestry (# Hets) | Genetic Ancestry (# Hets) |
|---|---|---|---|---|---|---|---|---|---|
| 17:43049191:TG:T | *BRCA1* | Frameshift | rs80357590 | c.5335delC | p.Gln1779fs | Pathogenic (3*) | 1 | ESA (1) | Filipino and Southeast Asian (1) |
| 17:43051071:A:C | *BRCA1* | Missense | rs41293463 | c.5324T>G | p.Met1775Arg | Pathogenic (3*) | 3 | AA (3) | African American and African Diaspora (3) |
| 17:43057062:T:TG | *BRCA1* | Frameshift | rs80357906 | c.5266dupC | p.Gln1756fs | Pathogenic (3*) | 6 | EA (6) | AJ (5), Non-AJ European (1) |
| 17:43063345:TTTTC:T | *BRCA1* | Frameshift | rs80357867 | c.5177_5180delGAAA | p.Arg1726fs | Pathogenic (3*) | 1 | AA (1) | |
| 17:43063903:G:T | *BRCA1* | Missense | rs28897696 | c.5123C>A | p.Ala1708Glu | Pathogenic (3*) | 1 | EA (1) | Non-AJ European (1) |
| 17:43063917:A:C | *BRCA1* | Stop gained | rs80356974 | c.5109T>G | p.Tyr1703Ter | Pathogenic (3*) | 1 | EA (1) | Non-AJ European (1) |
| 17:43071017:TC:T | *BRCA1* | Frameshift | | c.4896delG | p.Ser1633fs | NA | 1 | EA (1) | Non-AJ European (1) |
| 17:43074403:C:A | *BRCA1* | Stop gained | rs80357366 | c.4603G>T | p.Glu1535Ter | Pathogenic (3*) | 3 | AA (3) | African American and African Diaspora (2) |
| 17:43076559:GC:G | *BRCA1* | Frameshift | rs1064793951 | c.4412delG | p.Gly1471fs | Pathogenic (3*) | 1 | HA (1) | Colombian (2) |
| 17:43082434:G:A | *BRCA1* | Stop gained | rs41293455 | c.4327C>T | p.Arg1443Ter | Pathogenic (3*) | 1 | O (1) | Non-AJ European (1) |
| 17:43091443:G:C | *BRCA1* | Stop gained | rs398122680 | c.4088C>G | p.Ser1363Ter | Pathogenic (3*) | 1 | AA (1) | African American and African Diaspora (1) |
| 17:43091455:T:TGC | *BRCA1* | Frameshift | | c.4074_4075dupGC | p.Gln1359fs | NA | 1 | ESA (1) | Filipino and Southeast Asian (1) |
| 17:43091669:CCTCA:C | *BRCA1* | Frameshift | rs80357842 | c.3858_3861delTGAG | p.Ser1286fs | Pathogenic (3*) | 1 | HA (1) | Central and South American (1) |
| 17:43091714:G:A | *BRCA1* | Stop gained | rs80357208 | c.3817C>T | p.Gln1273Ter | Pathogenic (3*) | 1 | HA (1) | Puerto Rican (1) |
| 17:43091771:TAGAC:T | *BRCA1* | Frameshift | rs80357868 | c.3756_3759delGTCT | p.Ser1253fs | Pathogenic (3*) | 2 | EA (2) | Non-AJ European (1) |
| 17:43091924:G:A | *BRCA1* | Stop gained | rs62625308 | c.3607C>T | p.Arg1203Ter | Pathogenic (3*) | 2 | EA (1), HA (1) | Puerto Rican (1) |
| 17:43092119:C:A | *BRCA1* | Stop gained | rs886040126 | c.3412G>T | p.Gly1138Ter | Pathogenic (3*) | 1 | AA (1) | African American and African Diaspora (1) |
| 17:43092196:TCTTG:T | *BRCA1* | Frameshift | rs80357701 | c.3331_3334delCAAG | p.Gln1111fs | Pathogenic (3*) | 2 | AA (1), HA (1) | African American and African Diaspora (1), Central and South American (1) |
| 17:43092570:C:CT | *BRCA1* | Frameshift | rs886040088 | c.2960dupA | p.Ser988fs | Pathogenic (3*) | 1 | EA (1) | Non-AJ European (1) |
| 17:43092615:TC:T | *BRCA1* | Frameshift | rs80357573 | c.2915delG | p.Gly972fs | Pathogenic (3*) | 1 | AA (1) | African American and African Diaspora (1) |
| 17:43093055:TG:T | *BRCA1* | Frameshift | rs80357970 | c.2475delC | p.Asp825fs | Pathogenic (3*) | 1 | EA (1) | AJ (1) |
| 17:43093496:T:A | *BRCA1* | Stop gained | rs80357082 | c.2035A>T | p.Lys679Ter | Pathogenic (3*) | 1 | EA (1) | Non-AJ European (1) |
| 17:43093950:CTT:C | *BRCA1* | Frameshift | rs431825387 | c.1579_1580delAA | p.Lys527fs | Pathogenic (3*) | 1 | EA (1) | Non-AJ European (1) |
| 17:43094018:T:A | *BRCA1* | Stop gained | rs397508877 | c.1513A>T | p.Lys505Ter | Pathogenic (3*) | 1 | HA (1) | |
| 17:43094451:G:T | *BRCA1* | Stop gained | rs886037986 | c.1080C>A | p.Cys360Ter | Pathogenic (3*) | 1 | EA (1) | Non-AJ European (1) |
| 17:43094472:C:T | *BRCA1* | Stop gained | rs80356935 | c.1059G>A | p.Trp353Ter | Pathogenic (3*) | 1 | O (1) | Non-AJ European (1) |
| 17:43104260:A:C | *BRCA1* | Stop gained | rs80356936 | c.303T>G | p.Tyr101Ter | Pathogenic (3*) | 1 | AA (1) | African American and African Diaspora (1) |

| CHR:POS:REF:ALT | Gene | Function | rsID | cDNA Position | Protein Position | ClinVar Significance (Review Status) | # Hets | Self-Reported Ancestry (# Hets) | Genetic Ancestry (# Hets) |
|---|---|---|---|---|---|---|---|---|---|
| 17:43106457:T:C | *BRCA1* | Missense | rs80357382 | c.211A>G | p.Arg71Gly | Pathogenic (2*) | 3 | HA (2), ESA (1) | Puerto Rican (2), Filipino and Southeast Asian (1) |
| 17:43106478:A:C | *BRCA1* | Missense | rs80357064 | c.190T>G | p.Cys64Gly | Pathogenic (2*) | 1 | AA (1) | African American and African Diaspora (1) |
| 17:43106487:A:C | *BRCA1* | Missense | rs28897672 | c.181T>G | p.Cys61Gly | Pathogenic (3*) | 1 | EA (1) | Non-AJ European (1) |
| 17:43115744:C:T | *BRCA1* | Missense | rs80357498 | c.116G>A | p.Cys39Tyr | Pathogenic (2*) | 1 | M (1) | |
| 17:43124027:ACT:A | *BRCA1* | Frameshift | rs80357914 | c.68_69delAG | p.Glu23fs | Pathogenic (3*) | 41 | EA (36), M (4), O (1) | AJ (38) |
| 17:43124030:C:CT | *BRCA1* | Frameshift | rs80357783 | c.66dupA | p.Glu23fs | Pathogenic (3*) | 1 | O (1) | Non-AJ European (1) |
| 13:32319298:G:T | *BRCA2* | Stop gained | rs397507646 | c.289G>T | p.Glu97Ter | Pathogenic (3*) | 1 | O (1) | Non-AJ European (1) |
| 13:32329467:CTG:C | *BRCA2* | Frameshift | rs768580992 | c.658_659delGT | p.Val220fs | Pathogenic (3*) | 1 | EA (1) | Non-AJ European (1) |
| 13:32330986:TGACA:T | *BRCA2* | Frameshift | rs80359659 | c.755_758delACAG | p.Asp252fs | Pathogenic (3*) | 1 | EA (1) | Non-AJ European (1) |
| 13:32331009:C:T | *BRCA2* | Stop gained | rs80358998 | c.772C>T | p.Gln258Ter | Pathogenic (3*) | 2 | EA (2) | Non-AJ European (2) |
| 13:32332517:C:T | *BRCA2* | Stop gained | | c.1039C>T | p.Gln347Ter | NA | 1 | AA (1) | African American and African Diaspora (1) |
| 13:32332778:AAAAG:A | *BRCA2* | Frameshift | rs80359277 | c.1310_1313delAAGA | p.Lys437fs | Pathogenic (3*) | 1 | HA (1) | Ecuadorian (1) |
| 13:32332862:G:T | *BRCA2* | Stop gained | | c.1384G>T | p.Glu462Ter | NA | 1 | AA (1) | |
| 13:32333165:TG:T | *BRCA2* | Frameshift | rs876659278 | c.1689delG | p.Trp563fs | Pathogenic (3*) | 1 | EA (1) | Non-AJ European (1) |
| 13:32333228:AAAAAG:A | *BRCA2* | Frameshift | rs80359302 | c.1755_1759delGAAAA | p.Lys585fs | Pathogenic (3*) | 1 | EA (1) | Non-AJ European (1) |
| 13:32333282:G:T | *BRCA2* | Stop gained | | c.1804G>T | p.Gly602Ter | NA | 1 | HA (1) | Puerto Rican (1) |
| 13:32333363:CTTACATT:C | *BRCA2* | Frameshift | rs1555282146 | c.1887_1893delTACATTT | p.Thr630fs | Pathogenic (3*) | 1 | AA (1) | African American and African Diaspora (1) |
| 13:32336684:G:GA | *BRCA2* | Frameshift | rs80359328 | c.2330dupA | p.Asp777fs | Pathogenic (3*) | 1 | EA (1) | Non-AJ European (1) |
| 13:32336750:A:T | *BRCA2* | Stop gained | | c.2395A>T | p.Lys799Ter | NA | 1 | EA (1) | AJ (1) |
| 13:32336781:T:G | *BRCA2* | Stop gained | rs397507285 | c.2426T>G | p.Leu809Ter | Pathogenic (3*) | 1 | EA (1) | Non-AJ European (1) |
| 13:32337055:TC:T | *BRCA2* | Frameshift | rs397507637 | c.2701delC | p.Ala902fs | Pathogenic (3*) | 1 | HA (1) | Puerto Rican (1) |
| 13:32337160:TA:T | *BRCA2* | Frameshift | rs398122753 | c.2808del | p.Lys936fs | Pathogenic (3*) | 1 | AA (1) | African American and African Diaspora (1) |
| 13:32337160:TAAAC:T | *BRCA2* | Frameshift | rs80359351 | c.2808_2811delACAA | p.Ala938fs | Pathogenic (3*) | 1 | HA (1) | Central and South American (1) |
| 13:32337305:G:GA | *BRCA2* | Frameshift | rs80359365 | c.2957dupA | p.Asn986fs | Pathogenic (3*) | 1 | EA (1) | Non-AJ European (1) |
| 13:32337312:A:AG | *BRCA2* | Frameshift | rs1555282969 | c.2957_2958insG | p.Asn986fs | Pathogenic (3*) | 1 | AA (1) | African American and African Diaspora (1) |
| 13:32337952:CTG:C | *BRCA2* | Stop gained | rs80359391 | c.3599_3600delGT | p.Cys1200Terfs | Pathogenic (3*) | 1 | EA (1) | Non-AJ European (1) |
| 13:32338034:CTG:C | *BRCA2* | Frameshift | rs80359395 | c.3680_3681delTG | p.Leu1227fs | Pathogenic (3*) | 1 | AA (1) | African American and African Diaspora (1) |
| 13:32338069:GA:G | *BRCA2* | Frameshift | rs80359401 | c.3717delA | p.Lys1239fs | Pathogenic (3*) | 1 | EA (1) | Non-AJ European (1) |
| 13:32338075:GT:G | *BRCA2* | Frameshift | NA | c.3723delT | p.Phe1241fs | Pathogenic (3*) | 1 | EA (1) | AJ (1) |

| CHR:POS:REF:ALT | Gene | Function | rsID | cDNA Position | Protein Position | ClinVar Significance (Review Status) | # Hets | Self-Reported Ancestry (# Hets) | Genetic Ancestry (# Hets) |
|---|---|---|---|---|---|---|---|---|---|
| 13:32338200:CTG:C | *BRCA2* | Frameshift | rs80359405 | c.3847_3848delGT | p.Val1283fs | Pathogenic (3*) | 3 | EA (2), M (1) | Non-AJ European (2), African American and African Diaspora (1) |
| 13:32338212:AAAAT:A | *BRCA2* | Frameshift | rs80359410 | c.3860_3863delATAA | p.Asn1287fs | Pathogenic (3*) | 1 | O (1) | African American and African Diaspora (1) |
| 13:32338277:G:T | *BRCA2* | Stop gained | rs80358638 | c.3922G>T | p.Glu1308Ter | Pathogenic (3*) | 8 | HA (8) | Puerto Rican (7) |
| 13:32338565:TC:T | *BRCA2* | Stop gained | rs398122777 | c.4211delC | p.Ser1404Terfs | Pathogenic (3*) | 1 | AA (1) | African American and African Diaspora (1) |
| 13:32338783:CA:C | *BRCA2* | Frameshift | rs886040531 | c.4429delA | p.Ile1477fs | Pathogenic (3*) | 1 | EA (1) | Non-AJ European (1) |
| 13:32338824:TACTG:T | *BRCA2* | Frameshift | rs80359451 | c.4471_4474delCTGA | p.Leu1491fs | Pathogenic (3*) | 1 | AA (1) | African American and African Diaspora (1) |
| 13:32338968:CT:C | *BRCA2* | Frameshift | | c.4616delT | p.Leu1539fs | NA | 1 | HA (1) | Central and South American (1) |
| 13:32338981:GA:G | *BRCA2* | Frameshift | rs80359461 | c.4631delA | p.Asn1544fs | Pathogenic (3*) | 1 | O (1) | Filipino and South East Asian (1) |
| 13:32338986:A:AG | *BRCA2* | Frameshift | | c.4631_4632insG | p.Asn1544fs | NA | 1 | AA (1) | |
| 13:32339069:CCTG:GCAAAGACC | *BRCA2* | Frameshift | | c.4716_4717delinsAAAGACC | p.Cys1573fs | NA | 1 | O (1) | Non-AJ European (1) |
| 13:32339084:GA:G | *BRCA2* | Frameshift | rs397507740 | c.4731delA | p.Glu1577fs | Pathogenic (3*) | 1 | AA (1) | African American and African Diaspora (1) |
| 13:32339318:T:TA | *BRCA2* | Frameshift | rs398122789 | c.4964dupA | p.Tyr1655fs | Pathogenic (3*) | 1 | EA (1) | Non-AJ European (1) |
| 13:32339320:C:G | *BRCA2* | Stop gained | rs80358721 | c.4965C>G | p.Tyr1655Ter | Pathogenic (3*) | 1 | M (1) | |
| 13:32339394:CTG:C | *BRCA2* | Frameshift | rs80359478 | c.5042_5043delTG | p.Val1681fs | Pathogenic (3*) | 1 | EA (1) | Non-AJ European (1) |
| 13:32339421:C:CA | *BRCA2* | Frameshift | rs80359479 | c.5073dupA | p.Trp1692fs | Pathogenic (3*) | 1 | EA (1) | Non-AJ European (1) |
| 13:32339421:CA:C | *BRCA2* | Frameshift | rs80359479 | c.5073delA | p.Lys1691fs | Pathogenic (3*) | 1 | HA (1) | Ecuadorian (1) |
| 13:32339460:CAGAA:C | *BRCA2* | Frameshift | NA | c.5110_5113delAGAA | p.Arg1704fs | Pathogenic (3*) | 1 | EA (1) | Non-AJ European (1) |
| 13:32339489:G:T | *BRCA2* | Stop gained | NA | c.5134G>T | p.Gly1712Ter | Pathogenic (3*) | 1 | EA (1) | Non-AJ European (1) |
| 13:32339571:ATTTAAGT:A | *BRCA2* | Frameshift | rs80359496 | c.5217_5223delTTTAAGT | p.Tyr1739fs | Pathogenic (3*) | 1 | EA (1) | Non-AJ European (1) |
| 13:32339699:C:CA | *BRCA2* | Frameshift | rs80359507 | c.5351dupA | p.Asn1784fs | Pathogenic (3*) | 1 | M (1) | Non-AJ European (1) |
| 13:32339699:CAA:C | *BRCA2* | Frameshift | rs80359507 | c.5350_5351delAA | p.Asn1784fs | Pathogenic (3*) | 1 | EA (1) | Non-AJ European (1) |
| 13:32339928:CAATT:C | *BRCA2* | Frameshift | rs770318608 | c.5576_5579delTTAA | p.Ile1859fs | Pathogenic (3*) | 1 | ESA (1) | Filipino and Southeast Asian (1) |
| 13:32340128:C:T | *BRCA2* | Stop gained | rs80358806 | c.5773C>T | p.Gln1925Ter | Pathogenic (3*) | 1 | EA (1) | AJ (1) |
| 13:32340149:CAT:C | *BRCA2* | Frameshift | rs763890036 | c.5796_5797delTA | p.His1932fs | Pathogenic (3*) | 1 | EA (1) | Non-AJ European (1) |
| 13:32340151:TAACC:T | *BRCA2* | Frameshift | rs80359538 | c.5799_5802delCCAA | p.Asn1933fs | Pathogenic (3*) | 1 | HA (1) | Puerto Rican (1) |
| 13:32340202:TGTTA:T | *BRCA2* | Frameshift | rs80359543 | c.5851_5854delAGTT | p.Ser1951fs | Pathogenic (3*) | 2 | HA (2) | Dominican (2) |
| 13:32340212:G:T | *BRCA2* | Stop gained | rs80358814 | c.5857G>T | p.Glu1953Ter | Pathogenic (3*) | 1 | AA (1) | African American and African Diaspora (1) |
| 13:32340300:GT:G | *BRCA2* | Frameshift | rs80359550 | c.5946delT | p.Ser1982fs | Pathogenic (3*) | 34 | EA (30), M (1), O (2) | AJ (30), Non-AJ European (1) |

| CHR:POS:REF:ALT | Gene | Function | rsID | cDNA Position | Protein Position | ClinVar Significance (Review Status) | # Hets | Self-Reported Ancestry (# Hets) | Genetic Ancestry (# Hets) |
|---|---|---|---|---|---|---|---|---|---|
| 13:32340392:A:T | *BRCA2* | Stop gained | rs80358840 | c.6037A>T | p.Lys2013Ter | Pathogenic (3*) | 2 | EA (2) | Non-AJ European (2) |
| 13:32340629:CTT:C | *BRCA2* | Frameshift | rs11571658 | c.6275_6276delTT | p.Leu2092fs | Pathogenic (3*) | 1 | EA (1) | |
| 13:32340751:A:AT | *BRCA2* | Frameshift | rs431825342 | c.6397dupT | p.Ser2133fs | Pathogenic (3*) | 1 | O (1) | Non-AJ European (1) |
| 13:32340816:ATC:A | *BRCA2* | Frameshift | rs80359596 | c.6468_6469delTC | p.Gln2157fs | Pathogenic (3*) | 2 | HA (1), O (1) | Non-AJ European (1) |
| 13:32340836:GACAA:G | *BRCA2* | Frameshift | rs770263702 | c.6486_6489delACAA | p.Lys2162fs | Pathogenic (3*) | 1 | HA (1) | Central and South American (1) |
| 13:32340882:ACATT:A | *BRCA2* | Frameshift | rs397507865 | c.6531_6534delTCAT | p.Ile2177fs | Pathogenic (3*) | 1 | HA (1) | Central and South American (1) |
| 13:32340983:GAA:G | *BRCA2* | Frameshift | rs80359611 | c.6629_6630delAA | p.Glu2210fs | Pathogenic (3*) | 1 | HA (1) | Central and South American (1) |
| 13:32340998:TACTC:T | *BRCA2* | Frameshift | rs80359616 | c.6644_6647delACTC | p.Tyr2215fs | Pathogenic (3*) | 1 | HA (1) | African American and African Diaspora (1) |
| 13:32341035:C:CA | *BRCA2* | Frameshift | rs1555284790 | c.6681dupA | p.Val2228fs | Pathogenic (3*) | 1 | HA (1) | Puerto Rican (1) |
| 13:32354920:TTC:T | *BRCA2* | Frameshift | rs756538291 | c.7069_7070delCT | p.Leu2357fs | Pathogenic (3*) | 1 | O (1) | Non-AJ European (1) |
| 13:32355068:C:CT | *BRCA2* | Frameshift | rs876659345 | c.7218dupT | p.Val2407fs | Pathogenic (3*) | 1 | EA (1) | AJ (1) |
| 13:32355212:GA:G | *BRCA2* | Frameshift | rs80359646 | c.7360delA | p.Ile2454fs | Pathogenic (3*) | 1 | EA (1) | Non-AJ European (1) |
| 13:32355270:T:TTC | *BRCA2* | Frameshift | | c.7417_7418insTC | p.Cys2473fs | NA | 1 | ESA (1) | |
| 13:32356463:C:T | *BRCA2* | Stop gained | rs80358971 | c.7471C>T | p.Gln2491Ter | Pathogenic (3*) | 2 | AA (2) | African American and African Diaspora (2) |
| 13:32356472:C:T | *BRCA2* | Stop gained | rs80358972 | c.7480C>T | p.Arg2494Ter | Pathogenic (3*) | 2 | EA (1), HA (1) | Non-AJ European (1), Dominican (1) |
| 13:32356550:C:T | *BRCA2* | Stop gained | rs80358981 | c.7558C>T | p.Arg2520Ter | Pathogenic (3*) | 2 | AA (2) | African American and African Diaspora (1) |
| 13:32357794:CAG:C | *BRCA2* | Frameshift | rs80359672 | c.7673_7674delAG | p.Glu2558fs | Pathogenic (3*) | 1 | HA (1) | Filipino and Southeast Asian (1) |
| 13:32362626:GCCTTT:G | *BRCA2* | Frameshift | rs80359686 | c.7913_7917delTTCCT | p.Phe2638fs | Pathogenic (3*) | 1 | EA (1) | AJ (1) |
| 13:32363498:AC:A | *BRCA2* | Frameshift | rs80359705 | c.8297delC | p.Thr2766fs | Pathogenic (3*) | 1 | EA (1) | Non-AJ European (1) |
| 13:32370483:TTA:T | *BRCA2* | Frameshift | rs397507984 | c.8415_8416delAT | p.Leu2805fs | Pathogenic (3*) | 1 | AA (1) | African American and African Diaspora (1) |
| 13:32379800:G:A | *BRCA2* | Missense | rs80359152 | c.9004G>A | p.Glu3002Lys | Pathogenic/Likely pathogenic (2*) | 2 | HA (1), ESA (1) | Dominican (1), Filipino and Southeast Asian (1) |
| 13:32379913:G:A | *BRCA2* | Synonymous | rs28897756 | c.9117G>A | p.Pro3039= | Pathogenic (2*) | 1 | EA (1) | Non-AJ European (1) |
| 13:32380085:C:T | *BRCA2* | Stop gained | rs80359180 | c.9196C>T | p.Gln3066Ter | Pathogenic (3*) | 2 | AA (2) | African American and African Diaspora (2) |
| 13:32380135:G:GA | *BRCA2* | Frameshift | rs80359752 | c.9253dupA | p.Thr3085fs | Pathogenic (3*) | 3 | EA (2), AA (1) | AJ (1), Non-AJ European (2) |
| 13:32394726:C:G | *BRCA2* | Stop gained | rs80359200 | c.9294C>G | p.Tyr3098Ter | Pathogenic (3*) | 1 | AA (1) | African American and African Diaspora (1 |
| 13:32394814:C:T | *BRCA2* | Stop gained | rs80359212 | c.9382C>T | p.Arg3128Ter | Pathogenic (3*) | 1 | M (1) | Non-AJ European (1) |
| 13:32394842:C:CT | *BRCA2* | Frameshift | rs876659435 | c.9413dupT | p.Leu3138fs | Pathogenic (3*) | 1 | EA (1) | Non-AJ European (1) |
| 13:32394863:CTG:C | *BRCA2* | Frameshift | rs80359763 | c.9435_9436delGT | p.Ser3147fs | Pathogenic (3*) | 1 | EA (1) | Non-AJ European (1) |

**Abbreviations:** Hets, heterozygous carriers; AA, African ancestry; AJ, Ashkenazi Jewish; EA, European ancestry; ESA, East or Southeast Asian ancestry; HA, Hispanic/Latino ancestry; M, multiple ancestries; O, Other ancestry.
cDNA and protein position provided for *BRCA1* ENST00000357654 (NM_007294.3) and *BRCA2* ENST00000380152 (NM_000059.3); Human reference genome build 38 (GRCh38)
